# Marine sponges as Chloroflexi hot-spots: Genomic insights and high resolution visualization of an abundant and diverse symbiotic clade

**DOI:** 10.1101/328013

**Authors:** Kristina Bayer, Martin T. Jahn, Beate M. Slaby, Lucas Moitinho-Silva, Ute Hentschel

## Abstract

*Chloroflexi* represent a widespread, yet enigmatic bacterial phylum. Meta-and single cell genomics were performed to shed light on the functional gene repertoire of *Chloroflexi* symbionts from the HMA sponge *Aplysina aerophoba*. Eighteen draft genomes were reconstructed and placed into phylogenetic context of which six were investigated in detail. Common genomic features of *Chloroflexi* sponge symbionts were related to central energy and carbon converting pathways, amino acid and fatty acid metabolism and respiration. Clade specific metabolic features included a massively expanded genomic repertoire for carbohydrate degradation in Anaerolineae and Caldilineae genomes, and amino acid utilization as nutrient source by SAR202. While Anaerolineae and Caldilineae import cofactors and vitamins, SAR202 genomes harbor genes encoding for co-factor biosynthesis. A number of features relevant to symbiosis were further identified, including CRISPRs-Cas systems, eukaryote-like repeat proteins and secondary metabolite gene clusters. *Chloroflexi* symbionts were visualized in the sponge extracellular matrix at ultrastructural resolution by FISH-CLEM method. *Chloroflexi* cells were generally rod-shaped and about 1 μm in length, albeit displayed different and characteristic cellular morphotypes per each class. The extensive potential for carbohydrate degradation has been reported previously for *Ca*. Poribacteria and SAUL, typical symbionts of HMA sponges, and we propose here that HMA sponge symbionts collectively engage in degradation of dissolved organic matter, both labile and recalcitrant. Thus sponge microbes may not only provide nutrients to the sponge host, but also contribute to DOM re-cycling and primary productivity in reef ecosystems via a pathway termed the “sponge loop”.

## Introduction

Sponges (Porifera) represent one of the oldest, still extant animal phyla. Fossil evidence shows their existence in the Precambrian long before the radiation of all other animal phyla (1, 2). Nowadays, sponges are globally distributed in all aquatic habitats from warm tropical reefs to the cold deep sea and are even present in freshwater lakes and streams (3). Sponges are increasingly recognized as important components of marine environments, due to their immense filter-feeding capacities and consequent impacts upon coastal food webs and biogeochemical (e.g., carbon, nitrogen) cycles (4, 5). Many marine sponges contain dense and diverse microbial consortia within their extracellular matrix (mesohyl). To date, 41 bacterial phyla (among them many candidate phyla) have been recorded from sponges, with recent amplicon sequencing studies suggesting up to 14,000 operational taxonomic units (OTUs) per sponge individual (6, 7). Sponges also constitute one of the most abundant natural sources of secondary metabolites, which are of commercial interest for the development of pharmaceuticals and new drugs (8) and are often produced by the microbial symbionts (9, 10).

Sponges can be classified into the so-called “high microbial abundance sponges” (HMA) harboring dense and diverse microbial consortia within their mesohyl tissues, and the “low microbial abundance sponges” (LMA) containing microbial numbers in the order of those found in seawater (11–13). While HMA sponges are enriched in *Chloroflexi, Acidobacteria, Ca*. Poribacteria, the LMA sponges are dominated by *Gamma*- and *Betaproteobacteria* as well as *Cyanobacteria*. Differences have also been observed with respect to functional gene content (14) pumping rates (15), and exchange of carbon and nitrogen compounds (16). There is mounting evidence that HMA sponges are specialized to feed on DOM while the LMA sponges preferably feed on particulate organic matter (POM), (7, 17, 18). It is thus tempting to speculate that the symbiotic microbiota of HMA sponges is involved in DOM degradation, and indeed, the microbiomes analyzed so far encode a diverse repertoire for carbon metabolism pathways and transporters for low molecular weight compounds (10, 19–21). However, the precise fluxes and mechanisms how DOM and POM are taken up and processed within the sponge holobiont remain unknown.

In the present study, we focused our metagenomic analyses on *Chloroflexi* as abundant and characteristic, yet understudied members of HMA sponge microbiomes. The phylum *Chloroflexi* comprises taxonomically and physiologically highly diverse lineages that populate a wide range of habitats (22–25) including the deep sea (26), uranium-contaminated aquifers (27) and the human oral cavity and gut (28, 29). *Chloroflexi* metabolism is very diverse, ranging from anxoxygenic photosynthesis, obligate aerobic/anaerobic heterotrophs, thermophiles, halophiles, clades capable of reductive halogenation, and even predators with gliding motility. Because only few *Chloroflexi* lineages have been cultivated (30) and because draft genomes are limited in number (26, 31), the specific functions of *Chloroflexi* within the ecosystem context remain frequently unknown.

Members of *Chloroflexi* are members of HMA sponge microbiomes, with representatives of classes SAR202, Anaerolineae, and Caldilineae, being most abundant (32). Visualization of *Chloroflexi* by fluorescence *in situ* hybridization (FISH) revealed bright and abundant signals (33, 34). Because *Chloroflexi* likely play an important role in the HMA sponge holobiont, we aimed (i) to assess the relative abundances and distribution in diverse HMA sponge species by using the largest dataset currently available (EMP sponge microbiome), (ii) to provide an updated phylogenetic analysis, (iii) to characterize the functional gene repertoire with a particular focus on carbon degradation and symbiotic lifestyle, and, (iv) to visualize *Chloroflexi* in mesohyl tissues at ultrastructural resolution by FISH-CLEM methodology. We applied a broad range of state-of-the-art methods, from global sponge surveys to single cell genomics and microscopy, to acquire comprehensive insights into the lifestyle of *Chloroflexi* symbionts.

## Materials and methods

### Relative abundance of Chloroflexi in high microbial abundance (HMA) sponges

To investigate the abundance of the bacterial phylum *Chloroflexi* on a global scale, sponge species, microbiome data from HMA sponges, classified and predicted (Cluster 1), were obtained from Moitinho-Silva *et al.*(2017). This dataset is a rarefied operational taxonomic unit (OTU) abundance matrix (23,455) from the mothur processed data of the Sponge Microbiome Project (32). Abundance of *Chloroflexi* OTUs were grouped according to the class level based on SILVA taxonomy (36). Relative abundances were calculated and displayed using the superheat R package (37).

### Sponge sampling, cell separation and handling

The *Chloroflexi* bins from an *Aplysina aerophoba* microbiome derived Illumina data set had been generated in a previous study (21). Briefly, a sponge individual was collected from Piran, Slovenia (45.5099 N; 13.5600 E), microbial cells were enriched from sponge tissues by differential centrifugation, The metagenomic DNA was isolated (33), sequenced (Illumina HiSeq2000 platform, 150 bp paired-end reads), and quality filtered by DOE Joint Genome Institute (JGI). Sequence data of metagenome bins are available from IMG under the Gold study ID Gs0099546 (Table 1). Data normalization, assembly and binning were conducted as described previously [Illumina-only assembly and binning published in (21)].

**Table 1:**
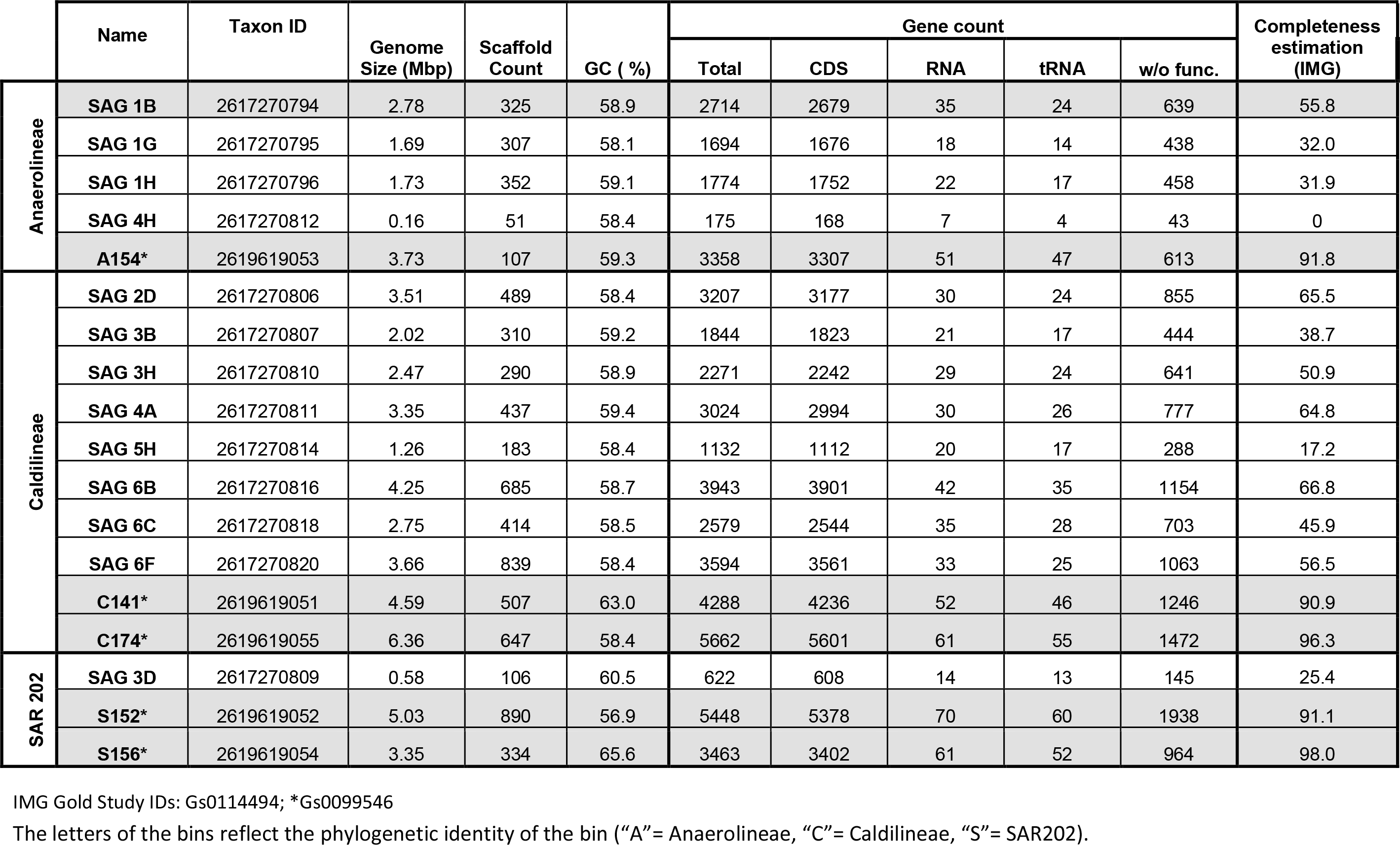
Overview of genomes analyzed in this study.

Single cells were sorted and their DNA amplified as described in Kamke *et al.* (19) and stored at −80°C in 96-well plates. Single amplified genomes (SAGs) were PCR screened using the universal primers 27f and 1492r to detect *Chloroflexi* 16S rRNA genes (38). SAGs tested positive for the presence of a single *Chloroflexi* 16S rRNA gene, were sequenced at GATC GmbH (Konstanz, Germany) on an Illumina MiSeq Personal Sequencer (300 bp; paired-end). Sequences were decontaminated using the IMG/EMR (Integrated microbial genomes & environmental samples) web tools following the single cell data decontamination protocol provided at JGI webpage (https://img.jgi.doe.gov/w/doc/SingleCellDataDecontamination.pdf). Contamination-free data of SAGs are available from IMG under the Gold study ID Gs0114494 (http://img.jgi.doe.gov/, for more details see Table 1).

### Phylogenetic tree construction

16S rRNA genes from one metagenome bin and all 13 single amplified genomes (SAGs) were manually quality checked and aligned with several reference sequences obtained from Silva database (SSU release 132) using the SINA aligner (39). The program MEGA 7.0.4 (40) was used for the protein alignment based on amino acid sequences of nine ribosomal proteins (L2, L4, L14, L15, L22, L24, S3, S17, S19), determination of best tree construction model (GTR+G+I model for 16S rRNA genes and JTT model for proteins), and final tree construction (Neighbor Joining method). As references for the protein tree, protein sequences from all public available *Chloroflexi* genomes were included (Suppl. table 1). Due to low genome completeness, some of the used proteins were missing in the regarding genomes. The trees were visualized using iTOL (interactive tree of life; http://www.itol.embl.de/).

### Fluorescence in situ hybridization and FISH-CLEM

FISH probes were designed based on the 16S rRNA gene alignment for sponge specific clades within the classes Anaerolineae and Caldilineae using the probe design tool implemented in ARB (41). Candidate probes were tested *in silico* for their specific hybridization conditions using different target and non-target reference sequences using mathFISH (http://mathfish.cee.wisc.edu/). Probes with best performance were tested for hybridization specificity on fixed (4% paraformaldehyde) *A. aerophoba* microbial cell preparations as described in Fieseler *et al.* (33) using formamide (FA) concentration gradients. Finally, we used for Caldilineae probe Cal825 (5’-[Cy3]-ACACCGCCCACACCTCGT-[Cy3]-3’, *E.coli* binding position: 825-843) and for Anaerolineae probe Ana1005 (5’-[Alexa647]-TCCGCTTTCGCTTCCGTA-[Alexa647]-3’, *E.coli* binding position: 1005-1023). Additionally, probe SAR202-104 (5’-[Alexa488]-GTTACTCAGCCGTCTGCC-[Alexa488]-3’, *E.coli* binding position: 104-122) was used to identify members of SAR202 group in sponges (42). All probes were double labelled at 5’-and 3’-ends (Sigma-Aldrich, Steinheim, Germany). To establish the newly designed sponge specific *Chloroflexi* probes and the previously published SAR202-104R probe for the sponge microbiome, FISH conditions were optimized using microbial cell preparations from *A. aerophoba*. The three probes did not co-localize using 10%, 20%, and 30% FA demonstrating specific binding of the probes to the *Chloroflexi* classes/ clades in standard FISH experiments (Figure S6).

For ultrastructural visualization of sponge *Chloroflexi*, we applied a recently established FISH-CLEM protocol (Fluorescence *in situ* Hybridization Correlated Light and Electron Microscopy (43). Briefly, freshly sampled *A. aerophoba* sponges were transported to the University of Wuerzburg where small mesohyl discs (2 mm diameter, 200 μm thickness) were subjected to high pressure freezing (HPF) and freeze substitution. Samples were embedded in LR white and 100 nm ultrathin sections were cut using a Histo Jumbo Diamond Knife (Diatome AG, Biel, Switzerland) on a Leica EM UC7 ultramicrotome (Leica Microsystems, Wetzlar, Germany). The sections were placed on poly-L-lysine coated slides and subjected to fluorescence *in situ* hybridization with the Chloroflexi clade specific probes at 10% FA concentration (900 mM NaCl, 20 mM Tris/HCL pH 7.4, 0.01% sodium dodecyl sulphate, 20% dextran sulfate). All three class or clade specific probes were co-hybridized and fluorescence signals were detected using an Axio Observer.Z1 microscope equipped with AxioCam 506 and Zen 2 version 2.0.0.0 software (Carl Zeiss Microscopy GmbH, Göttingen, Germany). On the same sections that were used for fluorescence microscopy, SEM was carried out using a field emission scanning electron microscope JSM-7500F (JEOL, Japan) with LABE detector (for back scattered electron imaging at extremely low acceleration voltages) directly on the microscope slides. FISH and SEM images of same regions were computer-correlated based on sponge heterochromatin pattern as described in Jahn *et al.* (43).

### Functional genomic analysis

Genomic data from single amplified genome sequences and the extracted metagenome bins were loaded and analyzed in IMG (http://img.jgi.doe.gov/) using the KEGG Orthology (KO) terms assigned to our data sets and metabolic pathways (KEGG) were analyzed. To identify CRISPR related genes, CRISPRfinder (http://crispr.u-psud.fr/Server/) was used. For the search of specific metabolite gene cluster, antiSMASH was used (44). The genomic potential of investigated microbial symbionts to degrade and transform complex carbohydrates was assessed by screening the IMG-predicted open reading frames (ORFs) of the genome data against the dbCAN (45) and classified according to the carbohydrate-active enzymes (CAZymes) database (46).

## Results and discussion

### Chloroflexi abundance in HMA sponges

Recently, members of the phylum *Chloroflexi* were shown to be present in much higher abundance and diversity in HMA sponges than in LMA sponges, which is why they were termed “indicator species” for HMA sponges (35). Here, we provide further details into the presence and abundance of *Chloroflexi* in sponges that were either classified or predicted by machine learning as HMA (35), (Figure 1, tables S2A and S2B). The recently compiled Sponge Microbiome Project (6, 32) was used as a reference database. In these 63 investigated sponge species, *Chloroflexi* abundances ranged from 4.39 % ± 3.02 % *(Chondrilla caribensis)* to 31.89 % ± 5.27 % *(Aplysina* sp.), (Figure 1 - right panel, suppl. Table 2A). With respect to the *Chloroflexi* classes, the SAR202 clade was the most abundant, contributing on average to 47.74% ± 22.00% of the phylum total abundance (Figure 1 - top panel, suppl. table 2B). Members of the classes Caldilineae (22.35 % ± 17.93%) and Anaerolineae 11.64% ± 12.30% were also abundant. Unclassified OTUs at class level represented 14.50% ± 10.77% of *Chloroflexi* sequences indicating that there is phylogenetic novelty still to be discovered. Despite some variability (Figure 1, main panel),the classes SAR202, Caldilineae and Anaerolineae as well as diverse hitherto unclassified OTUs dominated the *Chloroflexi* population in the HMA sponges. The remaining classes amounted to a total of 3.78% of total phylum abundance.

**Figure 1:**
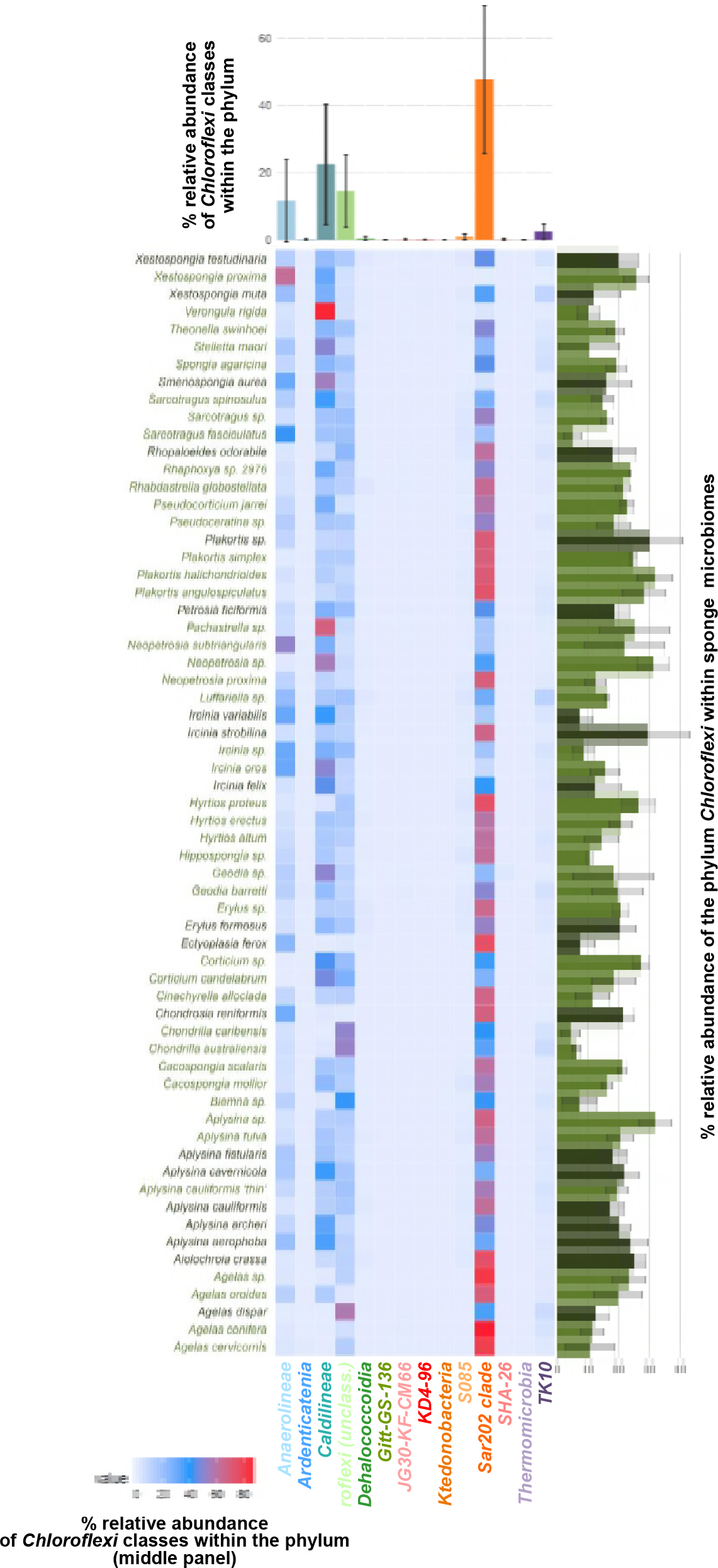
Relative abundance of *Chloroflexi* classes in HMA sponges extracted from Earth Microbiome Project (EMP) data (32). The central panel shows the mean relative abundance of classes per sponge species. Top panel shows mean relative abundance of *Chloroflexi* classes in all HMA sponges (± standard deviation). Right panel displays the mean relative abundance of the phylum *Chloroflexi* in predicted (light green bars) and classified (dark green bars) HMA sponges (± standard deviation).

**Table 2:** Sponge specific features of selected genomes.

| Class/ clade | Name | Taxon ID | CRISPR |  | ANK | secondary metabolite gene cluster |  |  |
| --- | --- | --- | --- | --- | --- | --- | --- | --- |
|  |  |  | CRISPRfinder<br>Total (No of repeats per spacer) | IMG<br>Total |  | type 1 PKS | terpene | other |
| Anaerolineae | SAG 1B | 2617270794 | - | - | 1 | - | - | - |
|  | A154* | 2619619053 | - | - | 2 | - | - | - |
| Caldilineae | C141* | 2619619051 | 7 (27, 24, 13, 7, 3, 30, 4) | 9 | 9 | 2 | - | - |
|  | C174* | 2619619055 | 5 (21, 24, 8, 32, 27) | 8 | 1 | - | 1 | - |
| Sar202 | S152* | 2619619052 | 5 (7, 34, 26, 7, 4) | 9 | 5 | 3 | 4 | 1 |
|  | S156* | 2619619054 | 1 (13) | 3 | 2 | 1 | 2 | 1 |

IMG Gold Study IDs: Gs0114494; \*Gs0099546 - extracted metagenome bins; ANK: Ankyrins and Ankyrin-repeat containing proteins; secondary metabolite cluster found using antiSMASH 3.0; values are total numbers of genes per genome. The letters of the bins reflect the phylogenetic identity of the bin ("A"= Anaerolineae, "C"= Caldilineae, "S"= SAR202).

### Phylogeny of Chloroflexi metagenome bins and single amplified genomes (SAGs)

16S rRNA gene sequencing of 260 single amplified genomes (SAGs) obtained from sponges resulted in the identification of 13 SAGs that belonged to the phylum *Chloroflexi*. Phylogenetic analysis of 16S rRNA genes revealed that one (3D), four (1B, 1G, 1H, 4H) and eight (2D, 3B, 3H, 4A, 5H, 6B, 6C, 6F) SAGs belonged to the classes SAR202, Anaerolineae and Caldilineae, respectively, forming a well-supported sequence cluster (bootstrap: 100) with other sponge derived sequences (Figure S1). The binning of metagenomic sequence data (21) resulted in an additional five high-quality bins (Table 1). The only metagenome bin containing a 16S rRNA gene (S156) belonged to SAR202. The SAR202 sequences formed a well-supported cluster (bootstrap value: 98) with other sponge derived 16S rRNA gene sequences (Figure S1). To elucidate the affiliation of the remaining four metagenome bins lacking 16S rRNA genes of appropriate length, a concatenated genome tree based on nine ribosomal genes was calculated (Figure 2A). One bin (A154) was affiliated to the class Anaerolineae, two metagenome bins were associated to the Caldilineae (C141 and C174), and two were associated to the SAR202 clade (S152 and S156) within the phylum *Chloroflexi*. The phylogenetic affiliation of metagenome bin S156 was congruent with the 16S rRNA gene analysis. SAGs were included in the protein based phylogenetic analysis when they encoded at least three of the nine ribosomal genes. Due to the lack of more complete reference genomes from SAR202 microorganisms, the most complete one (ca. 25%, SAR202 cluster bacterium sp. SCGC AAA240-N13, Gs0017605, (26) was included in this analysis although only one ribosomal protein could be used for tree construction. Both analyses showed a stable phylogeny of all SAGs and metagenome bins to above described classes or clades within the phylum *Chloroflexi*. All three classes/clades were visualized in the *A. aerophoba* sponge mesohyl matrix by fluorescence in situ co-hybridization (FISH) on ultra-thin tissue sections using class/ clade-specific probes. *Chloroflexi* cell signal was abundant, especially for SAR202 and cells were metabolically active as judged by the brightness of the FISH probe (Figure 2B).

**Figure 2A:**
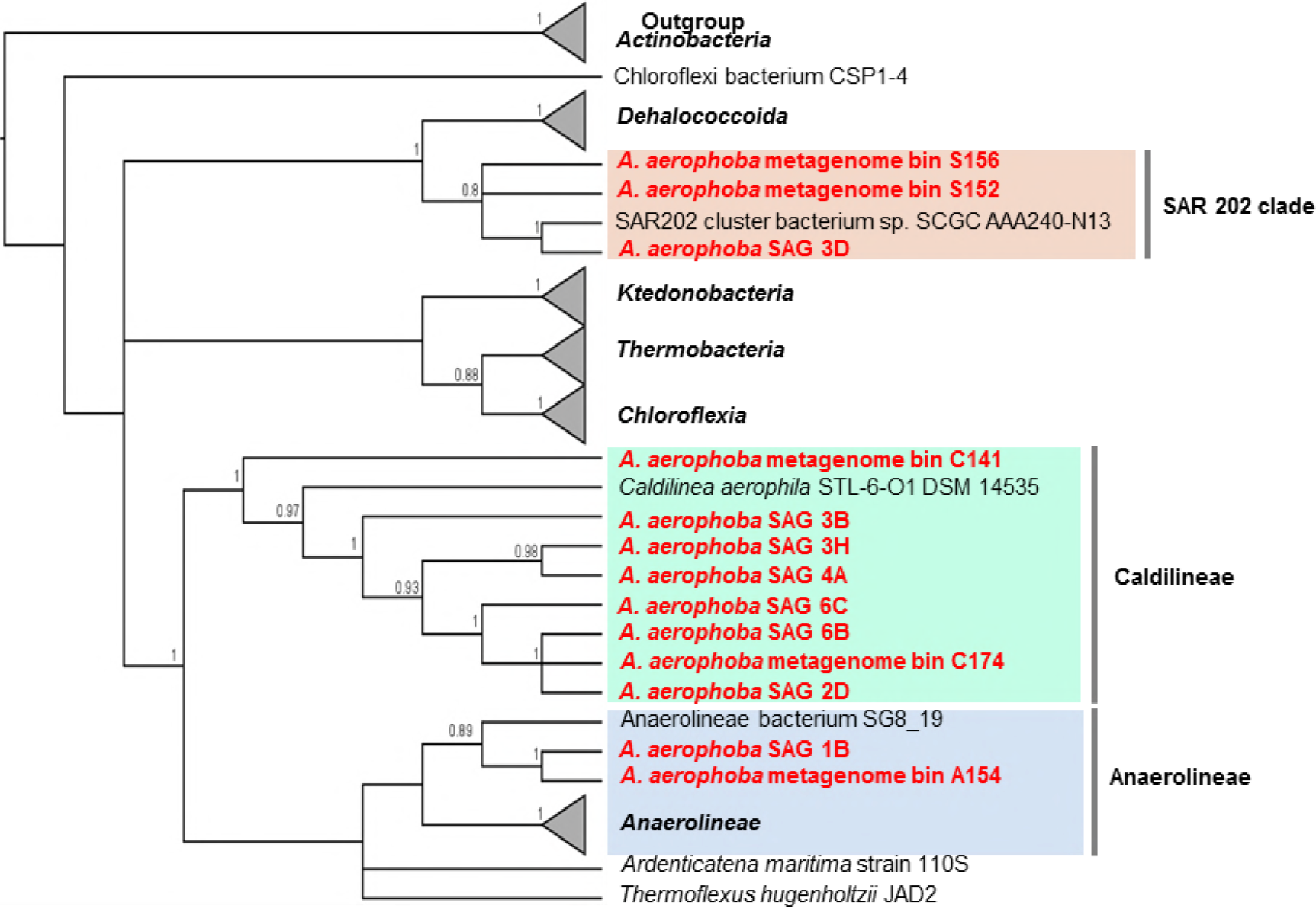
Concatenated protein tree. Maximum Likelihood phylogenetic analysis of *Chloroflexi* metagenome bins and SAGs (in red) derived from 1914 positions of 60 sequences. The percentage of replicate trees in which the associated taxa clustered together in the bootstrap test (100 replicates) are shown as symbols (biggest closed circles bs 100, symbol size represent values from 75 and above). Initial tree for the heuristic search were obtained automatically by applying Neighbor-Joining and BioNJ algorithms to a matrix of pairwise distances estimated using a JTT model. Used reference genomes with accession numbers can be found in Table S1.

**Figure 2B:**
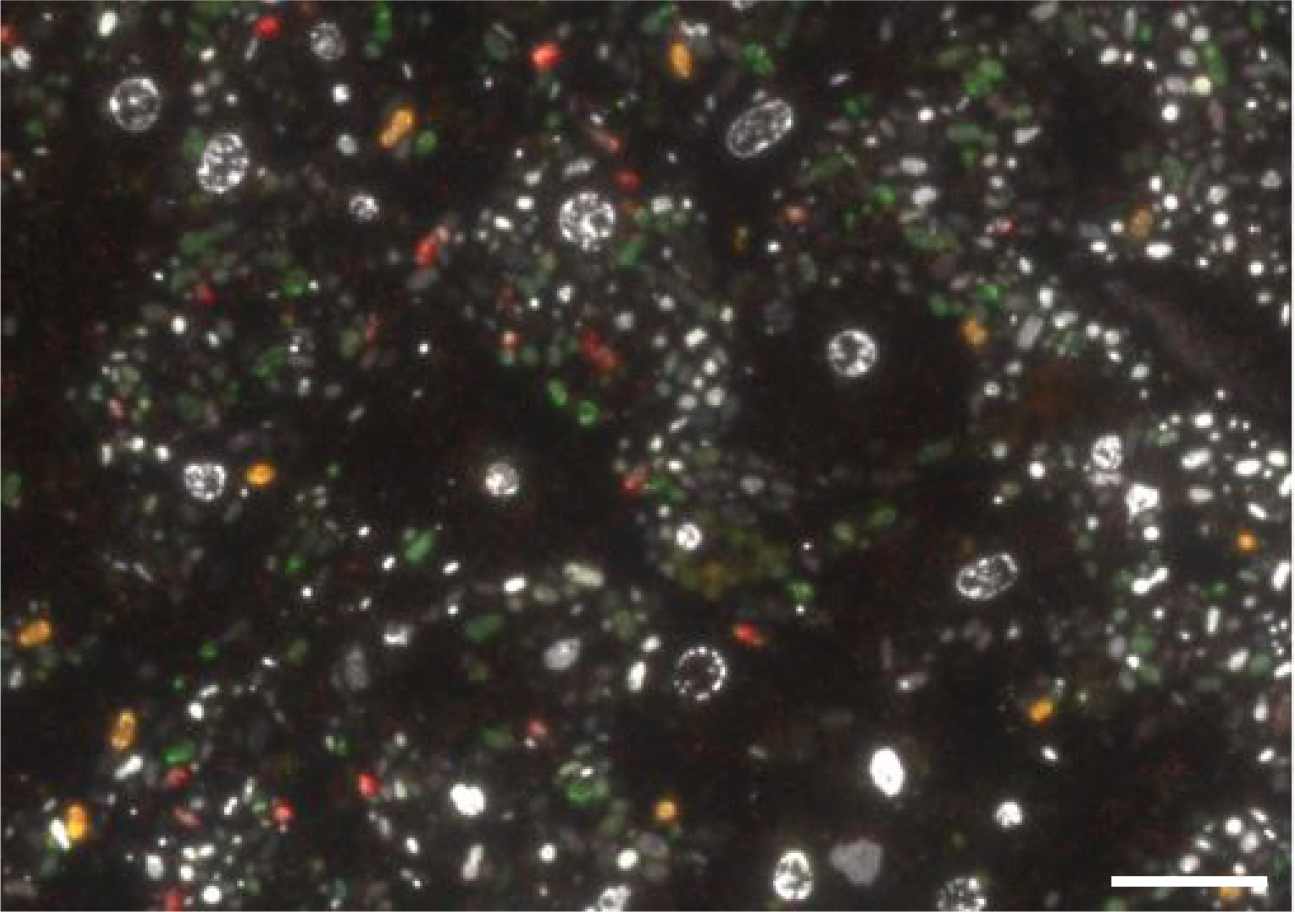
Distribution of *Chloroflexi* clades in *Aplysina aerophoba* mesohyl using fluorescence *in situ* hybridization (FISH): SAR202 cells are displayed in green, Caldilineae cells in orange, Anaerolineae cells in red. The nucleotide stain DAPI (white/ grey) served as reference for the localization of unstained cells. Scale bar: 10 μm.

### General description of genomes

Final genome assembly sizes for the sponge associated *Chloroflexi* single cells (SAGs) ranged from 0.16 to 4.25 Mbp, representing up to 66.85 % of genome completeness derived from IMG based estimations (Table 1). Final genome assembly sizes for the metagenomic bins ranged from 3.3 - 6.3 Mbp, showing high estimated genome completeness between 82.8 - 93.4% (Table 1). The guanine-cytosine (GC) content ranged from 58.08 to 59.32%, from 58.36 to 62.98% and from 56.93 to 65.59% for Anaerolineae, Caldilineae and SAR202, respectively. The number of identified genes was highly variable, ranging from 3,358 genes for Anaerolineae genome bin A154, to 5,448 for the SAR202 bin S152 and 5,662 genes for the Caldilineae bin C174 (Table 1). The five metagenome bins (two for Caldilineae (C141, C174), two for SAR202 (S152, S156) and one for Anaerolineae (A154)) which had > 90% coverage were chosen for detailed metabolic analysis and inner-phylum comparison. Due to the lack of marine reference genomes for Anaerolineae and Caldilineae or incompleteness of SAR202 reference genomes (26) we did not compare sponge derived Chloroflexi with these references. The letters of the bins were chosen to reflect their phylogenetic identity (“A”= Anaerolineae, “C”= Caldilineae, “S”= SAR202). A gene or enzyme is considered present when identified in both bins of the corresponding clade. Additionally, the most complete Anaerolineae SAG 1B (55.76% genome completeness estimation) was included in the analysis. For Anaerolineae, we consider an enzyme or gene present when identified in bin A154 and the SAG 1B was taken as additional support.

### Central metabolism of sponge associated Chloroflexi

Metabolic reconstruction suggests that *Chloroflexi* are aerobic and heterotrophic bacteria (see supporting text and supplementary figures for details). Genes involved in glycolysis and the tricarboxylic acid cycle (TCA) were almost completely identified in all metagenome bins (Figure S2A, B). The pentose phosphate pathway (PPP), including the oxidative and non-oxidative phase is largely present. Also, the Entner-Doudoroff-pathway was identified, but lacks the gene encoding for enzyme phosphogluconate dehydratase (EC: 4.2.1.12) in all clades. Furthermore, the enzyme 2-dehydro-3-deoxyphosphogluconate aldolase (EC: 4.2.1.14) is missing in SAR202 (Figure S2C). Interestingly, only the genomes of Anaerolineae and Caldilineae encode for enzymes involved in the ribulose monophosphate pathway (conversion of β-D-fructose-6P to D-ribulose-5P), which was originally found in methylotrophic bacteria but is now recognized as a widespread prokaryotic pathway involved in formaldehyde fixation and detoxification (47).

With respect to autotrophic carbon fixation, the reductive citrate acid cycle (Arnon-Buchanan cycle), is largely present, with the exception of ATP-citrate lyase (EC: 3.2.2.8), that is missing in all six genomes. A second pathway of autotrophic carbon fixation, the Wood-Ljungdahl-pathway was partially identified. While the genes encoding for carbon monoxide dehydrogenase (EC: 1.2.99.2) and formate dehydrogenase (EC: 1.2.1.43) are noticeably present in all six genomes, the rest of the Wood-Ljungdahl-pathway remains incomplete (see Figure S2D). Ammonia import and assimilation is encoded on all investigated genomes, but SAR202 and Caldilineae have additional genes for glutamate synthesis from glutamine and directly from ammonia. The transport of nitrite (and possibly also nitrate) is encoded on all investigated genomes while the reduction to ammonia is encoded only by SAR202 (Figure S2E). The incorporation of sulfur (with thiosulfate or few other sulfur compounds as donor) into S-containing amino acids might be possible in all clades whereas the assimilatory reduction of sulfate is restricted to Anaerolineae and Caldilineae genomes (Figure S2F).

Genes encoding for enzymes of the respiratory chain, including succinate dehydrogenase, cytochrome c oxidase, NADH dehydrogenase and an f-type ATPase, are largely represented on all genomes. These energy gaining processes additionally provide precursors for further metabolic pathways such as biosynthesis of purines and pyrimidines, amino acids and co-factors, or structural compounds. Machinery for transcription and translation, purine and pyridimidine metabolism are largely present. Fatty acid (FA) biosynthesis and degradation pathways were detected in all six genomes. Genes involved in FA beta-oxidation were found almost completely (supporting text), but also the three key enzymes involved in the propionyl-CoA pathway for odd-length and methylated fatty acid degradation were found among the genomes. This includes propionyl-CoA carboxylase (EC: 6.1.4.3) which was annotated in all six genomes, methylmalonyl-CoA epimerase (EC 5.1.99.1), and methylmalonyl-CoA mutase (EC: 5.4.99.2) both of which were found in all genomes except in S152. All genomes encode a number of different ABC transporters to supplement for nutrition and cell growth related compounds (incl. oligopeptides, phosphate, L-and branched chain amino acids, minerals as iron (III) and molybdate, metal ions as zinc, manganese and iron (II)). Additionally, all six genomes largely encode enzymes needed for biosynthesis of most amino acids (see supporting text). We could not identify any of the typical phosphotransferase systems, as it was the case for Poribacteria described previously (19).

We found genomic potential for aromatic degradation in *Chloroflexi* genomes, but pathways remain incomplete (supporting text). Several genes encoding for phenylpropionate and cinnamate degradation, terephthalate degradation, catechol degradation, and xylene degradation were identified on *Chloroflexi* genomes. Also, genes encoding for enzymes involved in ring-cleavage by Baeyer-Villinger oxidation and beta oxidation as well as ring-hydroxylating dioxygenases and isomerases were identified which could be involved in degradation of aromatic compounds. This finding is interesting in the context that many sponge species contain secondary metabolites that serve as a defense strategy against predators and biofouling (48). On the sponge genus level, highest (20-30% relative to the total microbiome) and most consistent presence of Chloroflexi within a sponge genus were found in the sponge genera Plakortis, Agelas (with the exception of *A. dispar*), *Aplysina* and sister taxon *Aiolochroia*. Interestingly, all of which contain characteristic natural products with aromatic ring structures that serve as chemotaxonomic markers (plakortolides, oroidins, bromo tyrosine alkaloids, respectively). It is therefore tempting to speculate that *Chloroflexi* and SAR202 presences and abundances are shaped, at least to some extent, by the natural products chemistry of their corresponding host sponges.

With respect to cell wall structure, the Anaerolineae and Caldilineae genomes encode the gene repertoire for peptidoglycan biosynthesis. The noticeable lack of peptidoglycan biosynthesis genes in the SAR202 genomes (supporting text) is consistent with previous analyses of three *Chloroflexi* genomes derived from uranium-contaminated aquifers (27). Synthesis pathways encoding for lipopolysaccharides or biosynthesis pathways for other glycan-based membranes could also not be annotated. The synthesis of an S-layer was proposed for SAR202 bacteria (26) as well as a member of GIF09 clade of *Chloroflexi* (27) but genes involved in sialic acid formation (N-Acetylneuraminic acid - Neu5Ac) in the amino and nucleotide sugar metabolism pathway are incomplete (supporting text). Nevertheless, the SAR202 bin S152 encodes a type 2-ABC transporter (NodJI) to export lipo-oligosaccharides. These compounds were shown to play a role in nodulation process in rhizobium bacteria (49), but their potential role for sponge-associated bacteria remains unclear. Additionally, consistent with previous observations (50) none of the six genomes encoded flagellar and chemotaxis genes.

### Metabolic specialization: extensive carbohydrate uptake and degradation in Anaerolineae and Caldilineae

The following features are metabolic specialities of Anaerolineae and Caldilineae and appear missing in SAR202, unless otherwise mentioned (Figure 3). We found a remarkable number of ABC transporters for the import of diverse mono- (ribose/ xylose, inositol, glycerol-3P and rhamnose) and oligosaccharides (sorbitol, raffinose/ stachyose/ melibiose, maltose, N-acetylglucosamine, arabinosaccharide) into Anaerolineae and Caldilineae cells. The pentoses xylose and ribose can also be imported into bacterial cells. Xylose can be processed to xylulose which may enter the pentose phosphate cycle finally leading into glycolysis. Ribose can be converted in 5-phosphoribosyl 1-pyrophosphate (PRPP) which is precursor for the biosynthesis of the amino acid histidine or it may fuel into purine and pyrimidine synthesis (supporting text). Both groups may also be able to import glycerol-3P which is a phosphoric ester of glycerol (a component of glycerophospholipids) which can be converted to fatty acids. Additionally, we found evidence for arabinose and rhamnose import and degradation; however annotation was incomplete (supporting text).

**Figure 3:** Summarized metabolic features which where found only in Anaerolineae and Caldilineae (left side, blue and green arrows) or in SAR202 genomes (right side, red arrows). The central metabolic pathways (glycolysis, TCA cycle, purine, pyrimidine histidine biosynthesis) located in the figure center are general features found in all genomes. Lines are dashed when pathways or transporter could not be annotated completely (single enzymes of the pathway or genes from the transporter where missing) or could not be annotated in both genomes of one clade.

Arabinooligosaccharides (such as α-L-arabinofuranosides, α-L-arabinans, arabinoxylans, and arabinogalactans) result from degradation of plant-like cell material entering the sponge by filtration. These substances may be imported by the almost completely annotated AraNPQ and MsmX transporters and be utilized to L-arabian and L-arabinose by the enzyme α-N-arabinofuranosidase (EC: 3.2.1.55, GH3). The enzyme L-arabinose isomerase (EC: 5.3.1.4, AraA) is present in all four genomes of Anaerolineae and Caldilineae and converts L-arabinose to L-ribulose which can further be converted by reactions of PPP to glucose-6P suitable for entering gylcolysis. Additionally, other oligosaccharides such as stachyose, raffinose, melibiose, and galactose can be imported and used in central metabolism (figure 3, supporting text).

The utilization of myo-inositol as carbon source and possibly as a regulatory agent was hypothesized previously for sponge-associated *Ca*. Poribacteria (19). Similarly, sponge-associated Anaerolineae and Caldilineae encode the nearly complete inositol degradation pathway (supporting text). Myo-inositol is likely degraded to glyceraldehyde-3-phosphate and acetyl-CoA, which are further used in the central metabolism. Inositol phosphates are found as part of eukaryotic and archaeal cell wall components (51). Phosphorylated inositol is a precursor for several lipid molecules including sphingolipids, ceramides and glycosylphosphatidylinositol anchors (52), as well as many stress-protective solutes of eukaryotes (51) and might be part of the signal transduction in sponges (53). Therefore, the sponge itself or eukaryotic microorganisms can probably provide inositol as a carbon source or regulatory agent for the microbial symbionts.

Uronic acids are sugar acids that can be found in biopolymers of plants, animals and bacteria (54, 55) and are known to occur in glycosaminoglycans (GAGs). GAGs in sponges are mainly composed of fucose, glucuronic acid (glucoronate), mannose, galactose, N-acetylglucosamine and sulfate (56–58). Enzymes involved in degradation of uronic acids were found in Anaerolineae and Caldilineae genomes. The possibility of galacturonate and glucuronate catabolism is supported by the conversion of 2-dehydro-3-deoxy-D-gluconate by enzymes glucoronate isomerase (EC: 5.3.1.12), tagaturonate reductase (EC: 1.1.1.58) and altronate hydrolase (EC: 4.2.1.7). Furthermore, the presence of genes encoding for oligogalacturonide lyase (EC: 4.2.2.6), 2-deoxy-D-gluconate 3-dehydrogenase (EC: 1.1.1.125) and 2-dehydro-3-desoxy-D-glucokinase (EC: 2.7.1.45) supports possible 4(4-α-D-gluc-4-enuronosyl)-D-galacturonate degradation activity. The products could then enter the ED pathway via 2-dehydro-3-desoxyphophogluconate aldolase (EC: 4.1.2.14). Uronic acid degradation could principally be connected to the inositol degradation pathway via D-galacturonate even though additional genome evidence, such as genes encoding for the enzyme inositol oxidase, (EC. 1.13.99.1) remain wanting (Figure 3, supporting text). A number of transporters for N-acetylglucosamine, digalacturonate, mannose and galactose (Figure 3, supporting text) were identified in Anaerolineae and Caldilineae genomes. Digalacturonate can be utilized by uronic acid degradation pathway (supporting text) and N-acetylglucosamine can be used directly in amino sugar and nucleotide sugar synthesis. The presence of uronic acid degradation pathways provides strong support that Anaerolineae and Caldilineae, similar to the previously described *Ca*. Poribacteria degrade glycosaminoglycan chains of proteoglycans, which are important components of the sponge host matrix (19). In that line, Anaerolineae and Caldilineae genomes were enriched in arylsulfatases A (Figure S3) which were discussed to be involved in metabolisation of sulfated polysaccharides from the sponge extracellular matrix (19, 21) and the heterotrophic ability of symbionts to use sponge components fur nutritional purposes.

### Expanded carbohydrate-active enzyme (CAZymes) repertoire in Caldilinae and Anaerolinae

In order to search for CAZymes, we screened the *Chloroflexi* genome data against dbCAN (45) and classified the enzymes according to the CAZy database (46). Most *Chloroflexi* hits were against glycosyl hydrolases (GH), glycosyl transferases (GT), and carbohydrate-binding modules (CBM). Consistent with the above described metabolic specializations, these enzyme classes were present in higher amounts in Caldilineae and Anaerolineae than in SAR202 (Figure S4). Altogether, 40 GH families were identified in all *Chloroflexi* genomes (Table S3). Glycosyl hydrolase family 109 was the most abundant family of GHs and was identified in all six genomes. GH109 family proteins are predicted as α-N-acetylgalactosaminidases (EC: 3.2.1.49) with putative substrates such as glycolipids, glycopeptides and glycoproteins all of which are common constituents of sponge mesohyl as well as dissolved organic matter from seawater. The family GH74 is second most abundant and is also present in all six genomes. These appear to be xyloglucan-hydrolyzing enzymes, that act on β-1,4 linkages and might help degrade various oligo- and polysaccharides. The previously reported glycosylhydrolases GH33 and GH32 (19, 20) were third most abundant, but were restricted to Caldilineae bin C174. This enzyme family is annotated as sialidase (EC: 3.2.1.18), capable of hydrolysing glycosidic linkages of terminal sialic acid residues, which are present in sponge mesohyl (59). Altogether 17 glycosyl transferases were identified on *Chloroflexi* genomes, with families GT2, GT4, and GT83 being most abundant. Among the 11 CBM families identified on *Chloroflexi* genomes, with CBM50 as the most abundant, but restricted to Caldilineae and Anaerolineae. CMB50 modules, also known as LysM domains, attach to various GH enzymes which are involved in the cleavage of chitin or peptidoglycan. The numbers of carbohydrate-active enzymes on *Chloroflexi* symbiont genomes reflect their extensive potential to degrade complex carbohydrates as was reported previously for *Ca*. Poribacteria and the sponge associated unidentified lineage SAUL (19, 20).

### Metabolic specialization: co-factor biosynthesis in SAR202 genomes

There is mounting evidence that vitamins and co-factors produced by diverse symbiont lineages could be beneficial to the sponge host (60–63). Parallel transcriptional activity profiling of the symbionts and the sponge showed that the symbionts had the capacity for vitamin B biosynthesis whereas the host transcripts displayed capacity for vitamin catabolism (64). It is thus tempting to speculate that the sponges’ nutrition is augmented by symbiont derived vitamins and cofactors. In the present study, at least two biosynthetic pathways for co-factor biosynthesis were identified on SAR202 genomes, which were absent in Anaerolineae and Caldilineae (Figure 4). Thiamine is an essential cofactor which is involved in central metabolism. The biosynthesis of the biologically active form thiamine diphosphate (TPP) from L-cysteine, glycine, pyruvate, and glyceraldehyde-3P is encoded on the SAR202 genomes (Figures 3, 4A). Although the pathway is incomplete, the data strongly suggest that the synthesis of TPP is restricted to SAR202 bacteria. Instead, Anaerolineae and Caldilineae appear to import thiamine via a ABC transporter (TbpA, ThiPQ) and convert it to TPP by use of thiamine pyrophosphokinase (EC: 2.7.6.2).

**Figure 4A:**
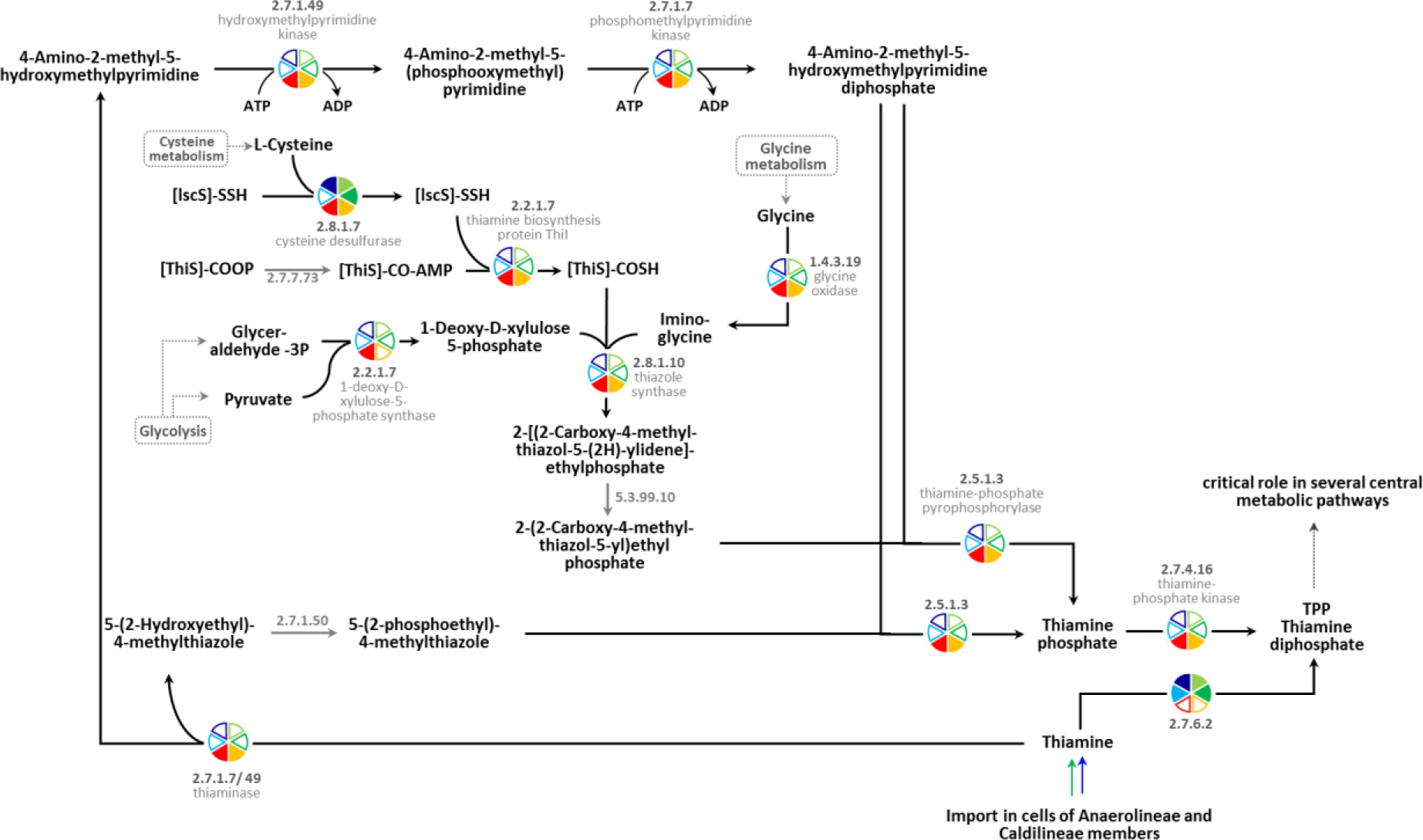
Pathway encoding for synthesis of thiamine in both SAR202 genomes (yellow and red pie symbols) could be annotated almost complete. Members of classes Anaerolineae and Caldilineae encode for the import of thiamine (blue and green arrow indicating import).

Secondly, riboflavin (vitamin B2) is required by enzymes and proteins to perform specific physiological functions. Specifically, the active forms, flavin mononucleotide (FMN) and flavin adenine dinucleotide (FAD), serve as cofactors for a variety of flavoprotein enzyme reactions. Most genomes encode for the enzymes FMN adenylyltransferase (EC: 2.7.7.2) and FAD riboflavin kinase (EC: 2.7.1.26) which activate riboflavin into FMN. However, only the SAR202 genomes encode the riboflavin biosynthesis enzymes which rely on GTP and ribulose-5P (Figures 3, 4B). Both substrates can be provided by pathways of the central metabolism pathways (purine metabolism and PPP).

**Figure 4B:** Pathway encoding for synthesis of riboflavin in both SAR202 genomes (yellow and red piesymbols) could be annotated almost complete. The conversion of Riboflavin in the biological active forms (FMN and FAD) was encoded in the genomes of all three classes.

### Potential for degradation of recalcitrant DOM in SAR202

The possible participation of deep sea SAR202 bacteria in degradation of recalcitrant or refractory DOM was recently postulated by Landry et al (26). Even though the exact composition of DOM in the world’s oceans remains to be elucidated, refractory DOM is an important component of the global carbon budget in terms of sheer mass. Landry *et al.* (26) argue that SAR202 genomes have an expanded repertoire of oxidative enzymes that may help in the oxidation of recalcitrant compounds. Interestingly, some of the major enzymes were also found to be enriched in SAR202 symbionts of sponges (Table S4). Among them are genes encoding proteins from the CaiB/BaiF family as well as related family III transferases. While the enrichment of CaiB was previously interpreted as carnitine being a carbon and nitrogen source for sponge symbionts (21), an alternative explanation may be that it serves to funnel substrates into degradation pathways without consumption of energy by shuffling CoA, thus generating free electrons (supporting text). Even though the precise function of CaiB/BaiF family proteins cannot be elucidated at the present time, the enrichment in SAR202 genomes is noteworthy. Further, a total of 53 flavin-dependent, class C oxidoreductases of the luciferase family (Flavin mononucleotide monooxygenases, FMNOs, COG2141) which include alkanesulfonate monooxygenase SsuD and methylene tetrahydromethanopterin reductase, were present and enriched in SAR202 genomes. These enzymes are proposed to participate in the oxidation of (long-chain) aldehydes to carboxylic acids, and/ or in cleavage of carbon-sulfur bonds in a variety of sulfonated alkanes (65). The SAR202 genomes contained 22 genes encoding for short-chain alcohol dehydrogenases (COG0300) which might be involved in canalization of ketone body derivate release. Some of these genes from bin S152 showed homologies to cyclopentanol and 3-α (or 20-β) - hydroxysteroid dehydrogenases which convert alicyclic-bound alcohol groups to ketones (26, 66). The combination of the enzymes described above could allow sponge associated *Chloroflexi* the conversion of recalcitrant alicyclic ring structure to more labile carboxylic acid, as proposed recently for SAR202 bacteria from deep sea (26).

Additionally, a number of oxidative enzymes was identified on the Chloroflexi genomes, but was not enriched in SAR202 (Table S4). These include a 2-oxoglutarate:ferrodoxin oxidoreductase (EC: 1.2.7.11) which oxidizes acetyl-CoA, carbon monoxide dehydrogenase (EC: 1.2.99.2) which might endow the bacteria to oxidize CO_2_ as described for some members of the Ktedonobacteria (67), CO- or xanthine dehydrogenases (COG1529) which are possibly involved in oxidation of a broad range of complex substrates (26), choline dehydrogenase (EC. 1.1.99.1) being possibly involved in the oxidation of alcohols to aldehydes, sarcosine oxidase (EC. 1.5.3.1), the serine hydroxymethyltransferase (2.1.2.1) with predicted function in choline degradation, formaldehyde dehydrogenase (EC: 1.2.1.46) and subunits of formate dehydrogenase (EC: 1.2.1.2/43) which oxidize formaldehyde and formate and might be involved in demethylation of various compounds. The overall presence and frequent enrichment of enzymes with oxidative capacity in SAR202 would be consistent with gene functions in degradation of recalcitrant DOM. However, owing to the sponges’ existence in shallow water sun-light benthic environments, it remains unclear whether the sponge symbionts encounter recalcitrant DOM derived from seawater sources. Alternatively, and similar to other high diversity microbiomes for example of ant, ruminant, and human guts (ref), the resident microbes are likely to specialize on certain substrates, thus promoting maximum nutrient exploitation and also securing their individual niche in the holobiont ecosystem.

### Symbiosis related features

Eukaryotic-like proteins (ELPs) seem to be a general genomic feature of sponge symbionts (20, 50, 60, 61, 68–70). Particularly ankyrin (ANK), tetratricopeptide (TPR), and leucine-rich (LRR) repeat proteins are postulated to be involved in mediating host-microbe interactions (71, 72). Ankyrin and ankyrin repeat containing proteins were detected in all six genomes (Figure S5, Table S5A). It was recently proposed that the expression of sponge symbiont derived ankyrin protein prevents phagocytosis by amoeba (73), and it is tempting to speculate that they protect the symbionts from digestion by the sponge archaeocytes in vivo. TPRs, possibly functioning as module for protein-protein interaction involved in a variety of cellular functions, including those that participate in bacterial pathogenesis (74) were found in all six genomes. However, LRRs were only identified in Caldilineae and SAR202 genomes (Figure S5, Table S5A). Many LRR proteins are involved in protein-ligand interactions; these include plant immune response and the mammalian innate immune response (for review see (75), such as the detection of pathogen-associated molecular patterns by recognition receptors (76). Our findings are in good agreement with general patterns previously found in metagenomes of sponge symbionts (50, 61), in enriched (mini)-metagenomes of cyanobacterial sponge symbionts (69) and single amplified genomes from members of SAUL (20) and *Ca*. Poribacteria (68).

Another example of sponge-symbiont enriched features are the clustered, regularly interspaced, short, palindromic repeats (CRISPRs) and their associated proteins (Cas) that have recently been reported from the genomes of sponge symbionts (20, 50, 60, 61, 69). Here, the investigated Caldilineae genomes and the two SAR202 genomes showed an enrichment of CRISPR (Table 2). However, the two most complete Anaerolineae genomes did not contain any CRISPR-Cas systems. The presence CRISPRs can be explained the extensive filter feeding activity of sponge hosts that result in high exposure of sponge symbionts and phages and other sources of free DNA from ambient seawater.

The synthesis of secondary metabolites is an important defense mechanism of sessile microorganisms such as sponges to protect against predators or biofouling (48). Many of these compounds are in fact produced by the sponge microbiome (10, 48) Especially polyketide synthases (PKS), non-ribosomal peptide synthetases (NRPS), and halogenases are regularly enriched in sponge symbionts, often with new structures and putatively novel activities (38, 61, 77–81). Here, we assessed the genomic repertoire of sponge associated Chloroflexi for secondary metabolism using antiSMASH (44). In both SAR202 genomes and in Caldilineae C141 we found up to three polyketide synthase (PKS) gene clusters all of which showed homologies to the previously reported type I PKS gene cluster from other sponge symbionts. Additional gene clusters for the production of terpenes and other yet to be identified substances were identified in the two SAR202 genomes and in Caldilineae C174 (Table 2, Table S5B). Both Anaerolineae genomes did not contain any gene clusters for the biosynthesis of secondary metabolites. While the exact functions of these gene clusters putatively involved in defense remain unknown, it appears that at least SAR202 bacteria and Caldilineae have the genomic repertoire for chemical defense within the sponge holobiont.

### Ultrastructural identification of sponge specific Chloroflexi

The correlation of probe specific fluorescence with scanning electron microscopy images (SEM) allowed the taxon-specific identification of *Chloroflexi* cells at ultrastructural resolution. Overall, distributions of all three *Chloroflexi* clades in *Aplysina aerophoba* indicate that SAR202 cells were more abundant than the other two *Chloroflexi* classes (Figure 5). This is consistent with the relative abundances of *Chloroflexi* classes in HMA sponges extracted from EMP data (Figure 1). Cells belonging to the SAR 202 clade (green signals in Fig. 5A and 5B) are generally rod shaped (0.8 × 1-2 μm) with a regular distribution of cell cytosol content. The Anaerolineae-specific probe targeted rod shaped cells (0.8 × 2.0μm) (red signal, Figure 5A). The characteristic feature of Caldilineae positive cells (ca. 1 × 2 μm) was the presence of electron dense capsules or mucus-like structures located at the cell poles (orange signal, Fig. 5B). All cells which were stained positive with a corresponding FISH probe showed a consistent morphology, which was taken as a measure of probe specificity.

**Figure 5:**
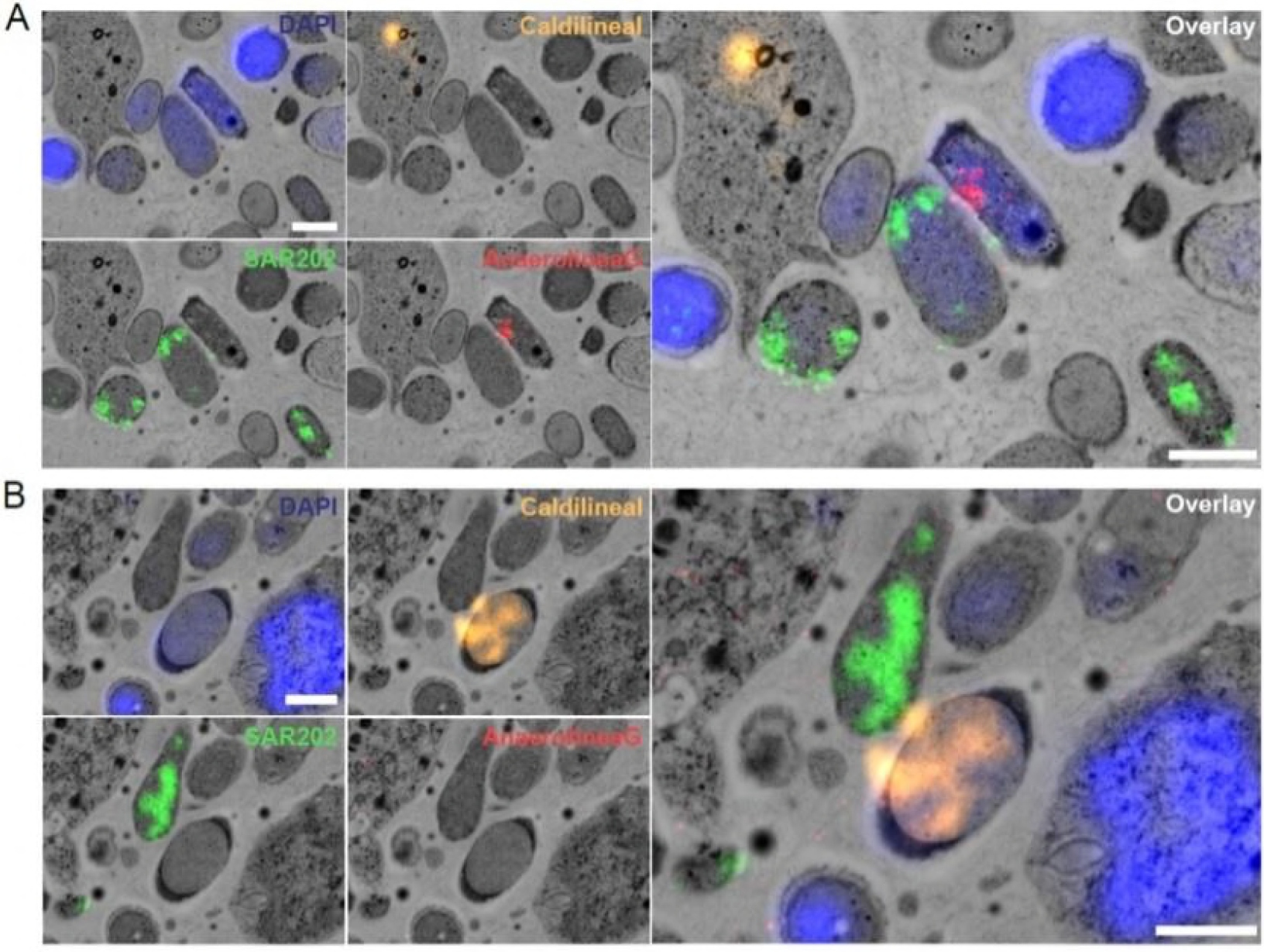
Visualization of sponge-associated *Chloroflexi* in *Aplysina aerophoba* mesohyl using FISH-CLEM: SAR202 cells are displayed in green (panel A and B) Caldilineae in orange (panel B) Anaerolineae in red (panel A). The nucleotide stain DAPI (blue) served as reference for the localization of unstained cells. In both panels the right picture is the overlay of all probes. Scale bars: 1 μm.

## Conclusions

Owing to the lack of cultivation and difficult experimental access for the majority of *Chloroflexi* clades, advancing knowledge has been limited to few lineages (26, 27, 31). The present study provides a new experimental opportunity as HMA sponges were identified as true *Chloroflexi* hotspots, both in terms of biomass and biodiversity. Meta-and single cell genomic analyses revealed metabolic specialization in that Anaerolineae and Caldilineae have an expanded gene repertoire for carbohydrate degradation while SAR202 specializes on amino acids. Similarly, while Anaerolineae/ Caldilineae take up cofactors, SAR202 has the genomic repertoire for their synthesis. A combination of FISH-CLEM allowed, for the first time, to visualize *Chloroflexi* in the host context and to identify characteristic cellular morphotypes. The results of this study suggest that *Chloroflexi* symbionts have the genomic potential for DOM degradation from seawater, both labile and recalcitrant. These findings are in line with previous reports that have shown extensive carbohydrate degradation potential in other HMA sponge symbionts (10, 19, 20). Thus, we hypothesize that collectively sponge microbes not only provide nutrients to the HMA sponge host, but also contribute to DOM cycling and primary productivity in reef ecosystems via a pathway termed the “sponge loop”. Considering the abundance and dominance of sponges in many benthic environments, we predict that role of sponge symbionts in biogeochemical cycles is larger than previously thought.

## Acknowledgements

We gratefully acknowledge financial support from the DFG (CRC1182-TPB1) and from the European Union’s Horizon 2020 research and innovation program under Grant Agreement No. 679849 (’SponGES’). B.M.S. and M.T.J. were supported by grants of the German Excellence Initiative to the Graduate School of Life Sciences, University of Wuerzburg. We thank Laura Rix and Lucia Pita Galan for many insightful discussions on sponge ecology and Christian Stigloher at University of Wuerzburg for support in FISH-CLEM.

## Figure legend

Table 1: Genomic features overview of single amplified genomes (SAGs) and metagenome bins of *A. aerophoba* associated *Chloroflexi* and closest relative reference genomes analyzed in this study.

Table 2: Genomic characteristics of the six genomes investigated in detail. The absolute numbers of CRISPR arrays - defined by CRISPRfinder and IMG, the number of Ankyrin repeat containing Proteins (ANK), as well as the number of secondary metabolite (antiSMASH) gene cluster per genome are shown.

Figure 1: Relative abundance of *Chloroflexi* classes in HMA sponges extracted from Earth Microbiome Project (EMP) data (6). The central panel shows the mean relative abundance of *Chloroflexi* classes per sponge species. Top panel shows mean relative abundance of *Chloroflexi* classes in all HMA sponges. Right panel displays the mean relative abundance of whole phylum *Chloroflexi* in HMA sponges (predicted and classified).

Figure 2A: Concatenated protein tree. Maximum Likelihood phylogenetic analysis of *Chloroflexi* metagenome bins and SAGs (in red) derived from 1914 positions of 60 sequences. The percentage of replicate trees in which the associated taxa clustered together in the bootstrap test (100 replicates) are shown as symbols (biggest closed circles bs 100, symbol size represent values from 75 and above). Initial tree for the heuristic search were obtained automatically by applying Neighbor-Joining and BioNJ algorithms to a matrix of pairwise distances estimated using a JTT model.

Figure 2B: Distribution of *Chloroflexi* clades in *Aplysina aerophoba* mesohyl using fluorescence *in situ* hybridization (FISH): The picture show the overlay of all probes, SAR202 cells are displayed in green, Caldilineae cells in orange, Anaerolineae cells in red. The nucleotide stain DAPI (white/ grey) served as reference for the localization of unstained cells. Scale bar: 10 μm.

Figure 3: Reconstruction of class-specific metabolic features. On the left side (blueish color) the transporter and utilization pathways found mainly in Anaerolineae and Caldilineae genomes are summarized. SAR202-specific transporter and pathways (for co-factor biosynthesis) are displayed on the right (reddish color). Common genomic features (glycolysis, amino acid metabolism and transport, fatty acid metabolism, respiratory chain etc.) are not shown.

Figure 4: Metabolic specialization in SAR202: Biosynthesis pathways of co-factors thiamine (A) and riboflavin (B) in SAR202 genomes.

Figure 5: Visualization of sponge-associated *Chloroflexi* in *Aplysina aerophoba* mesohyl using FISH-CLEM technique: SAR202 cells are displayed in green (panel A and B) Caldilineae in orange (panel B) Anaerolineae in red (panel A). The nucleotide stain DAPI (blue) served as reference for the localization of unstained cells. In both panels the right picture is the overlay of all probes. Scale bars: 1 μm.

